# PhageAI: a new approach to predicting the lifestyle of bacteriophages using proteinBERT and convolutional neural networks

**DOI:** 10.1101/2025.09.02.673651

**Authors:** Maria Urbanowicz, Arkadiusz Guziński, Żaneta Szulc

## Abstract

Bacteriophages are viruses that infect bacteria, including temperate, virulent and chronic phages. In the current times of increasing resistance to antibiotics (AMR), it is necessary to quickly find phages and determine their lifecycle, therefore computational methods in the first phase of composing phage preparations are necessary to find such sequences. The challenge in building such tools is the lack of good quality and correctly labeled data from multiple reference databases, which are necessary to select sequences to create cocktails that should contain complete and replication-capable phages. Current published models also do not support chronic phages, which constitute a small but significant group of phages. Another problem is explainability, current models are black-box models where we cannot observe what actually influences the model’s predictions, this may result in the model overlooking features that are truly associated with the phage life cycle. In the presented tool, however, these issues have been explicitly addressed. The presented tool was also compared to other lifecycle prediction models: Phatyp, PhagePred, Bacphlip and Phacts.

## Introduction

Phages are organisms that infect bacteria. Bacteriophages exhibit three main types of lifecycles. One of them is the temperate lifecycle, in which phages infect bacterial cells and integrate into their genome, remaining in a dormant state with the potential to enter the lytic cycle under favorable conditions (Mirzaei et al., 2017). In contrast, virulent phages exclusively follow the lytic cycle, rapidly lysing the host cell upon infection and releasing progeny virions (Shkoporov et al., 2018), while chronic phages are bacteriophage that replicates within the host cell without causing its lysis. New virions are released gradually leading to a persistent infection of the bacterium (Hobbs et al., 2016). Apart from the classical virulent, chronic and temperate phages, bacteriophages may also undergo pseudolysogeny, in which the phage genome remains in the cell in the carrier state, representing a stable coexistence of the virus with a bacterial population. Knowledge about bacteriophages lifecycles is continuously evolving (Chevallereau & Westra, 2021).

Bacteriophages are increasingly being considered a last-resort therapeutic option for individuals with antibiotic-resistant bacterial infections (Zalewska-Piątek, 2023). They are also applied in targeted therapies as vectors for genetic material (Hosseinidoust, 2017) and as modifiers of the intestinal microbiota (Mirzaei et al., 2022). In addition, their use in animal husbandry has been explored as an alternative or adjunct to antibiotics (Kahn et al., 2019).

High costs of their isolation, cultivation and observation, while the costs of Next Generation Sequencing (NGS) are decreasing, especially in the case of phages from metagenomic samples, means that a rapid method of verifying their life cycle is needed to be able to quickly haracterize them and determine potential practical application (Knezevic et al., 2021).

Currently, many attempts have been made to obtain a quick method of classification of bacteriophages. Among the available methods, we can mention PHACTS (McNair et al., 2012), a classifier that uses a random forest algorithm based on proteomic data, Bacphlip (Hockenberry et al., 2021), which also uses a random forest algorithm but is based on proteins described in the literature as being associated with the temperate cycle. Other classifiers are PhagePred (Song, 2020), which works based on Euclidean distance, using a Markov model based on k-mer frequency. There is also PhaTYP, which uses BERT as an algorithm that detects the patterns associated with a given phage life cycle based on the presence of phage protein clusters in sequences (Shang et al., 2023).

In this paper we present a novel, machine learning approach to classify bacteriophages into three groups: chronic, virulent and temperate on the basis of their proteomes. The novelty of the tool stems from its ability to incorporate the chronic cycle, explainability in the temperate virulent classification and usage of large protein embedding models to improve protein representation as described in detail in the Methods section. Our tool is available online at https://app.phage.ai/.

## Methods

### Dataset

For the training and testing sets, we used a dataset of 1,851 phages. To ensure representation of all possible genera, we selected representatives from 1,652 genera included in the Virus Metadata Resource (VMR) file (August 2022) from International Committee on Taxonomy of Viruses (ICTV), and we also included representatives for chronic phages. The chronic bacteriophages dataset also includes sequences derived from pig fecal samples (Billaud et al., 2021). Subsequently, observations similar more than 70 % were removed from the collection using the VIRIDIC program (Moraru et al., 2020) to ensure diversity and representativeness for each genus. Threshold 70% was established based on separation criteria of taxonomic groups for tailed phages (Turner et al., 2021).

### Preprocessing

The pre-filtered sequences were structurally annotated using the Glimmer program (Hyatt et al., 2017). The lifecycle labels for chronic phages were determined in case of morphogenesis protein (pI) or VP3 protein presence. Lifecycle labels for temperate and virulent phages were determined in the presence or absence of proteins associated with lysogenic lifecycle including integrase protein. All observations with labels are available in Supplementary Data (Table S3). The protein detection was performed using diamond (Buchfink et al., 2021), hmmscan (Eddy, 2011) and interproscan (Blum et al., 2021) using default parameters.

### Model preparation

For building model proteins sequences were used. Embedding was performed using the ProteinBertT5 model (Nadav et al., 2022), which was used as the first layer of the neural network model as embedding layer in the convolutional neural network (CNN) model. Each protein was first represented as an embedding vector. For each phage, a sequence was then constructed in which every protein corresponds to a ‘word’ represented by its ProtBertT5 embedding. These sequences were subsequently used as inputs for the model. The procedure diagram is available below (Fig. 1). Phage classification in our solution is based on two independent models. The first step is to classify bacteriophages into chronic and non-chronic phages, and the next step is to classify them into virulent and temperate phages (Fig. 2). For training both models, training and test sets were prepared independently. In each model, a random split of 80:20 was considered, taking into account the proportions of both classes for training models.

**Figure 1.**
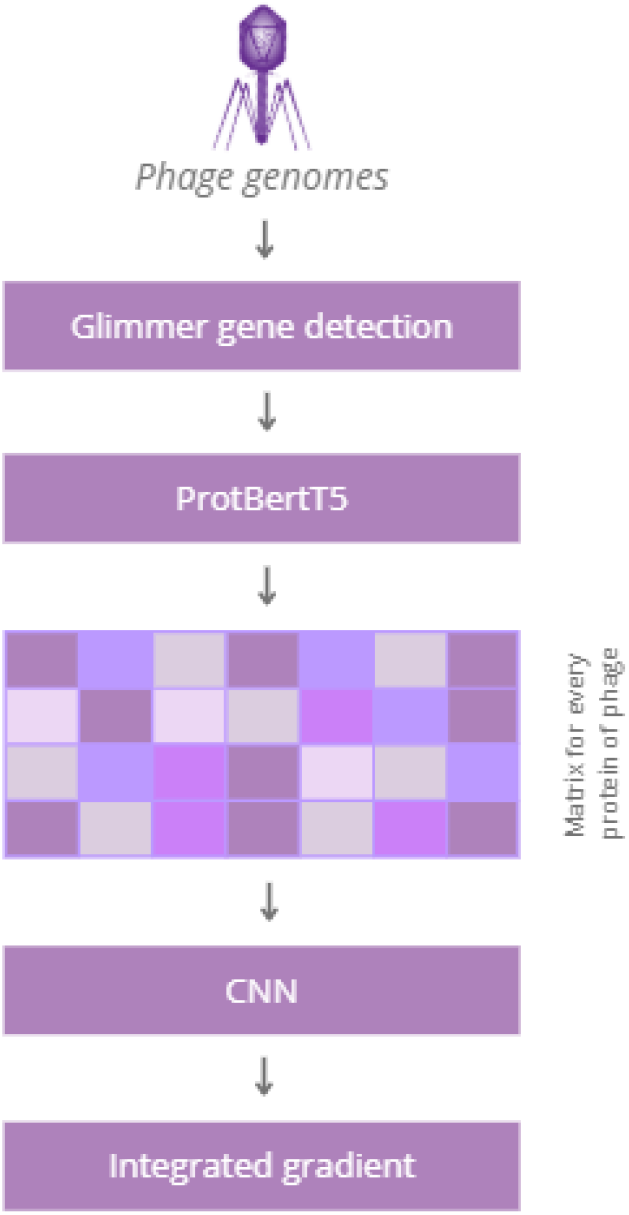
The technical model pipeline: genes were first detected using Glimmer, after which embeddings were generated for all bacteriophage proteins. A feature matrix was then constructed for classification, and model explainability was applied to distinguish virulent from temperate phages.

**Figure 2.**
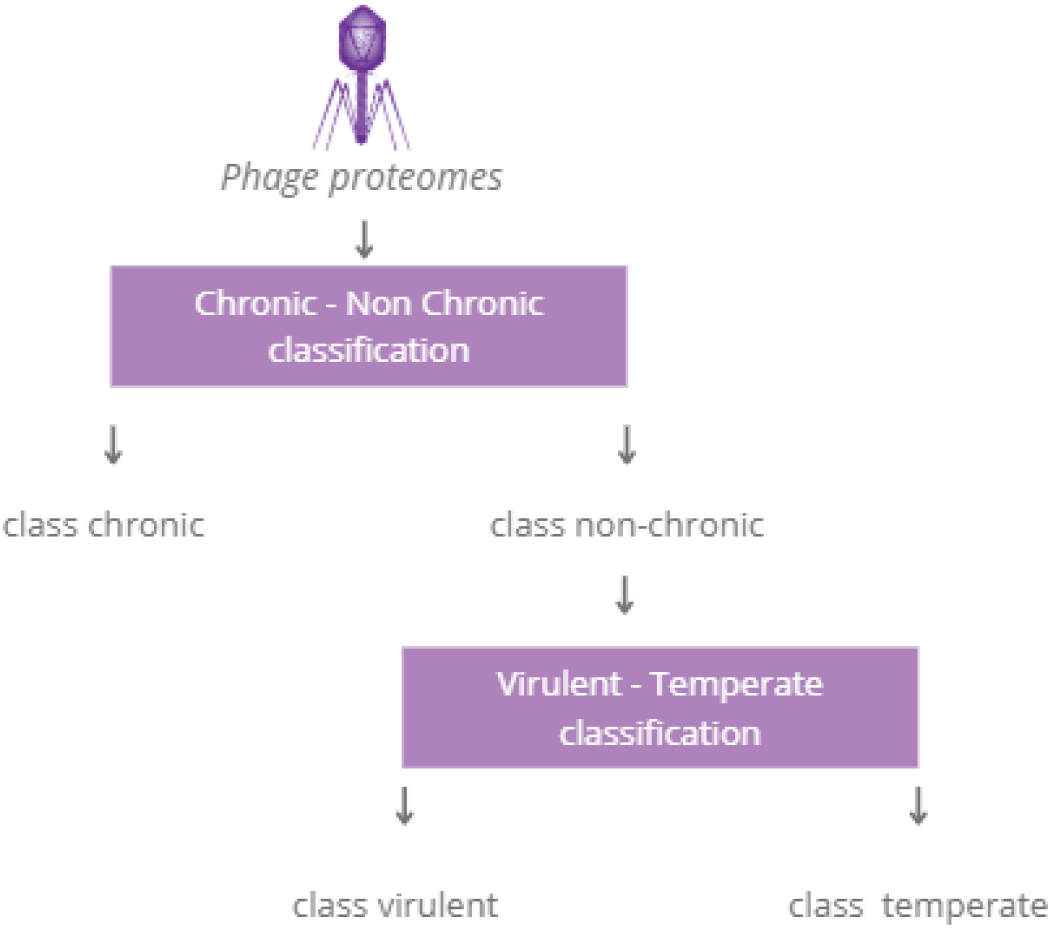
The PhageAI pipeline: the proposed methodology for bacteriophage life cycle recognition involves an initial classification of chronic versus non-chronic phages, followed by a subsequent classification into virulent or temperate classes.

### Explainability analysis

The integrated gradient (IG) method (Sundararajan et al., 2017) was used for explainability analyses, then the results from the IG analysis were combined using annotation from the diamond (Buchfink et al., 2021), hmmscan (Eddy, 2011) and interproscan (Blum et al., 2021) using default parameters programs to obtain a label for selected key proteins in the analysis. The annotation process was identical to the description of life cycle sets.

## Results

The results of the model were evaluated on the basis of an independent test set, for this purpose the following statistics were used: Precision (TP/TP+FP), Recall (TP/TP+FN)and F1-score (2 × (Precision × Recall) / (Precision + Recall)) and ROC-AUC curve. The original results were obtained on the test set of the classifier by comparing it with the available tools: Bacphlip, PhagePred, PhaTYP, Phacts.

The ROC curve analysis indicated that the best overall results were achieved using Bacphlip and PhageAI model. The results are also confirmed by precision, recall and F1-score statistics. Results for this comparison are available in Supplementary Data (Table S1). The added value of the publication is the result of explainability analyses for temperate and virulent bacteriophages, where proteins annotated as the integrases stand for 90% of the observations as the most important feature supporting predictions of temperate lifecycle, other proteins that were important for explainability analysis are listed in Supplementary Data (Table S2) for every bacteriophage from test set. Additionally, the table and figure contain results for chronic class, where evaluation on the test set demonstrates that the model attains highly satisfactory results across all performance metrics. (Fig. 3).

**Figure 3.**
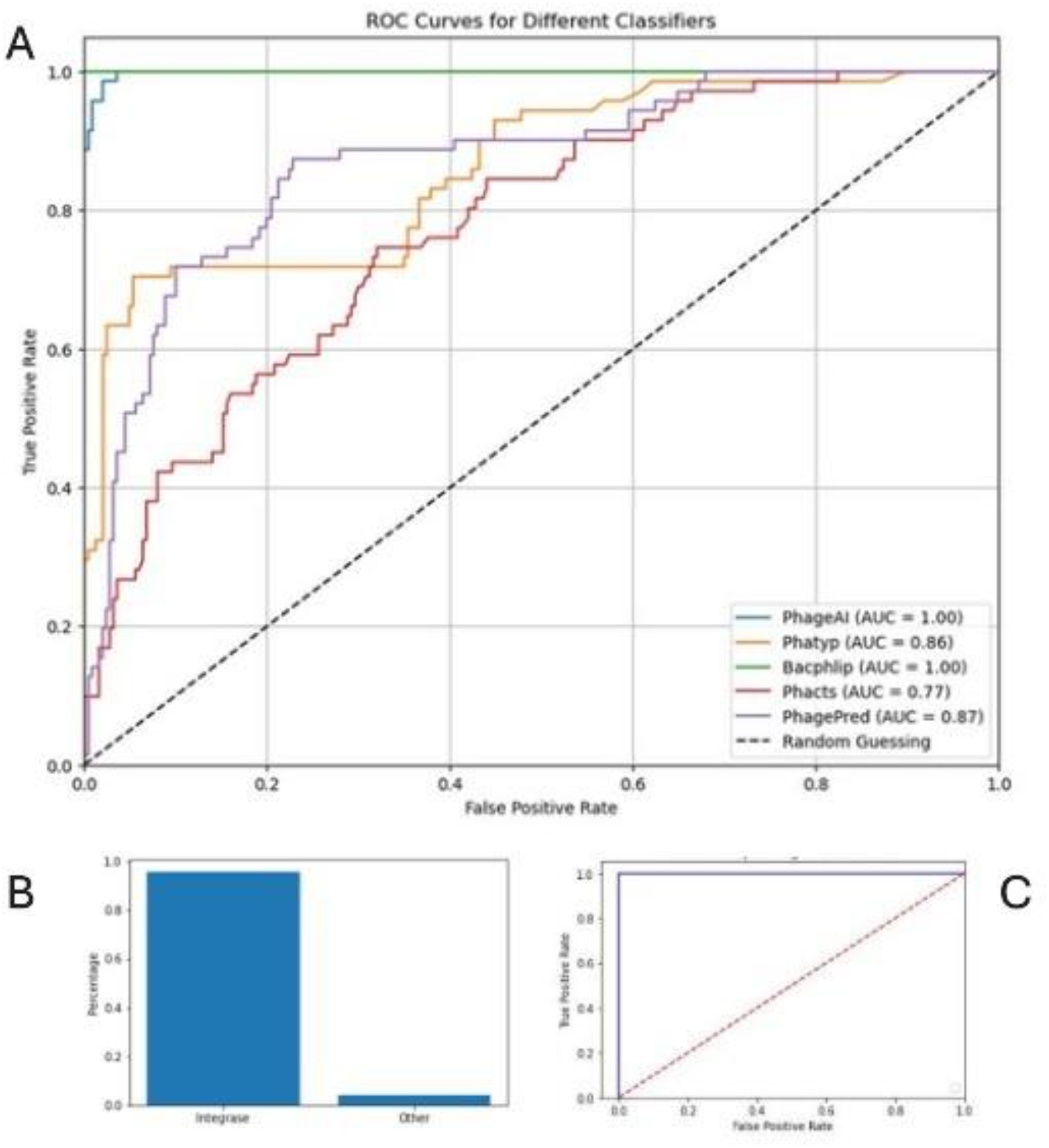
The ROC curve comparison on the test set. The value shown in the legend is AUCROC score. PhageAI and Bacphlip has the best performance (A). Proportion of the most important features for classification of temperate bacteriophages (B). The ROC curve for chronic-non chronic classificatory (C).

**Table 1.** The performance comparison on the test set.

|  | **Precision** | **Recall** | **F1-score** | **Metric** |
| --- | --- | --- | --- | --- |
| **PhageAI** | 0,99 | 0,99 | 0,99 | Virulent |
|  | 0,97 | 0,96 | 0,96 | Temperate |
|  | 0,98 | 0,98 | 0,98 | Weighted average |
| **PhaTYP** | 0,91 | 0,95 | 0,93 | Virulent |
|  | 0,79 | 0,68 | 0,73 | Temperate |
|  | 0,88 | 0,89 | 0,88 | Weighted average |
| **Bacphlip** | 0,99 | 0,98 | 0,99 | Virulent |
|  | 0,94 | 0,96 | 0,95 | Temperate |
|  | 0,98 | 0,98 | 0,98 | Weighted average |
| **PhagePred** | 0,95 | 0,65 | 0,77 | Virulent |
|  | 0,42 | 0,89 | 0,57 | Temperate |
|  | 0,84 | 0,70 | 0,73 | Weighted average |
| **Phacts** | 0,94 | 0,45 | 0,61 | Virulent |
|  | 0,32 | 0,90 | 0,47 | Temperate |
|  | 0,80 | 0,55 | 0,58 | Weighted average |
| **PhageAI for chronic phages** | 1,00 | 0,98 | 0,99 | Chronic |
|  | 1,00 | 1,00 | 1,00 | Non-chronic |
|  | 1,00 | 1,00 | 1,00 | Weighted average |

## Discussion

In an era of increased access to sequencing technology and a noticeable increase in the number of phage sequences published in databases (Fig. 4), the use of predictive models to answer one of the key questions: “What life cycle does a bacteriophage lead?” is beginning to gain importance.

**Figure 4.**
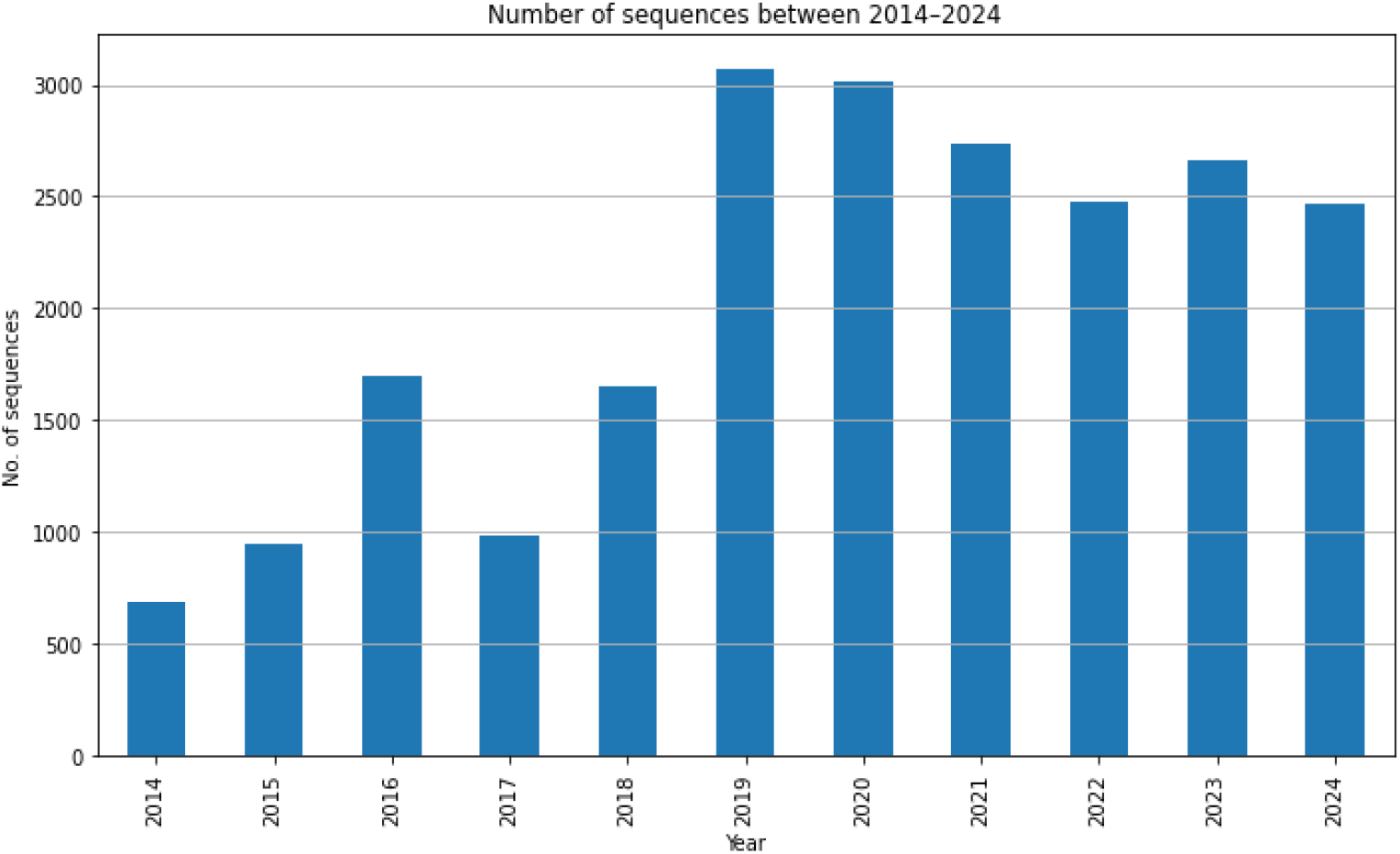
The number of phage sequences published in GenBank between 2014 and 2024. The figure shows an increase in the number of sequences since 2019, when the number exceeded 2,000 per year and has remained above that level since then.

Through our work, we wanted to address the opportunities offered by introducing AI/ML to phage science and the benefits it can bring. Working with models requires a large, well-characterized dataset. For this reason, we created the PhageAI service and adapted our bioinformatics pipelines to create the first datasets that could be used to create predictive models. We have achieved high levels of confidence and model explainability, something that is new for AI-based solutions and phage science. Our models utilized validated data and complete bacteriophages, focusing on nearly all currently described genera, ensuring the greatest possible diversity. This also allowed us to determine phage life cycles in taxonomic groups that were previously poorly described.

When comparing our model to existing approaches used in benchmarking, the model most closely aligned with our results is Bacphlip. This similarity is likely due to Bacphlip’s utilization of protein-level information and accurate annotation, which facilitates a better understanding of the factors influencing predictions. Properly annotated training data ensures that the assigned labels are accurate, improving model performance. A limitation of Bacphlip, however, is its inability to support chronic phages.

Interestingly, comparison with Phatyp yielded suboptimal results on our dataset. This may be attributed to its training on publicly available databases without prior validation, potentially leading to lower predictive accuracy on curated data. PhagePred and Phacts, being older models, likely show reduced performance due to reliance on outdated datasets and limited knowledge regarding bacteriophage lifecycle annotation at the time of their development.

Notably, none of the published models support chronic phages, despite the availability of current datasets that allow for the construction of representative training sets. Another challenge is the lack of model explainability, which prevents us from understanding the underlying basis of predictions. This limitation, especially when training and testing datasets are highly similar, can lead to potential bias or overestimation of model performance.

One of the limitations we see in presented model is the need to switch to amino acid sequences and the need to use additional programs for sequence annotation, which in itself may introduce errors in the form of predicting on incorrectly annotated proteins. On the other hand, by training the model on proteins, we managed to introduce methods of explaining the model’s indications that are understandable to humans. As our analyses show, the main determinant is the presence of integrase on the bacteriophage genome for most bacteriophages and those are results which met our expectations.

Our next step will be to configure an input data pipeline, which will allow us to ensure early on that the model is making predictions based on clean, complete phage sequences. Currently, the model will attempt to process a sequence regardless of whether it comes from a phage, bacteria, or other organism, which can lead to misunderstandings of results.

We anticipate that our work will facilitate research by accelerating bioinformatics analyses and making these approaches more accessible to researchers without extensive expertise in bioinformatics.

## Declarations

### Ethics approval and consent to participate

Not applicable.

### Consent for publication

Not applicable.

### Data availability

The dataset information can be found in **Supplementary Data.xlsx**

